# DEEPScreen: High Performance Drug-Target Interaction Prediction with Convolutional Neural Networks Using 2-D Structural Compound Representations

**DOI:** 10.1101/491365

**Authors:** Ahmet Sureyya Rifaioglu, Volkan Atalay, Maria Jesus Martin, Rengul Cetin-Atalay, Tunca Doğan

## Abstract

The identification of physical interactions between drug candidate chemical substances and target biomolecules is an important step in the process of drug discovery, where the standard procedure is the systematic screening of chemical compounds against pre-selected target proteins. However, experimental screening procedures are expensive and time consuming, therefore, it is not possible to carry out comprehensive tests. Within the last decade, computational approaches have been developed with the objective of aiding experimental studies by predicting novel drug-target interactions (DTI), via the construction and application of statistical models. In this study, we propose a large-scale DTI interaction prediction system, DEEPScreen, for early stage drug discovery, using convolutional deep neural networks. One of the main advantages of DEEPScreen is employing readily available simple 2-D images of compounds at the input level instead of engineered complex feature vectors that displayed limited performance in DTI prediction tasks previously. DEEPScreen learns complex features inherently from the 2-D molecular representations, thus producing highly accurate predictions. DEEPScreen system was trained for 704 target proteins (using ChEMBL curated bioactivity data) and finalized with rigorous hyper-parameter optimization tests. We compared the performance of DEEPScreen against shallow classifiers such as the random forest, logistic regression and support vector machines, to indicate the effectiveness of the proposed deep learning approach. Additionally, we compared DEEPScreen with other deep learning based state-of-the-art DTI predictors on widely used benchmark datasets and showed that DEEPScreen produces better or comparable results to the top performers. The method proposed here can be employed to computationally scan a large portion of the recorded drug candidate compound and protein spaces to aid the experimentalists working in the field of drug discovery and repurposing by providing a preselection of interesting novel DTIs.

## Introduction

One of the initial steps of drug discovery is the identification of novel drug-like compounds that interact with the predefined target proteins. *In vitro* and high-throughput screening experiments are performed to detect novel compounds with the desired interactive properties. However, high costs and temporal requirements makes it infeasible to scan massive target and compound spaces^1^. Due to this reason, the rate of the identification of novel drugs has substantially been decreased^2^. Currently, there are more than 90 million drug candidate compound records in compound and bioactivity databases such as ChEMBL^3^ and PubChem^4^, whereas, the size estimation for the whole “drug-like” chemical space is around 10^60^^5^. On the other hand, the current number of drugs (FDA approved or at the experimental stage) is around 10,000, according to DrugBank^6^. In addition, out of the 20,000 proteins in the human proteome, less than 3,000 of them are targeted by known drugs^7,8^. As the statistics indicates, the current knowledge about the drug-target space is limited, and novel approaches are required to widen our knowledge. Starting from the 2000s, developing computational methods to aid the drug discovery process by predicting the unknown interactions between drugs / drug candidate compounds and target biomolecules (i.e., drug target interaction -DTI-prediction or virtual screening)^9,10^ started to become a main-stream research area. Most of the DTI prediction methodologies are based on the idea that similar structures have similar activities^11,12^ and utilize the experimentally identified drug-target interaction information coming from bioassay results. DTI prediction methods usually employ supervised machine learning (ML) models to find new compounds (or targets) that possess features similar to the known drugs/targets^10,13,14^.

In DTA prediction methods, feature vectors correspond to fixed-dimensional quantitative representations/descriptors of the input samples (drugs and/or targets), used to characterize the molecular properties that play role in the interactions, so that the machine learning algorithm can learn from these features to accurately predict unknown DTIs. One of the essential steps in ML method development is the feature engineering, which constitutes designing, pre-processing and extracting meaningful features to be used for system training. In computational drug discovery studies, feature engineering is generally performed using computationally intensive third party methods/tools, where the main limitation is the constructed features not generalizing well to the whole proteochemical space^15^, also, they often suffer from the curse of dimensionality^16^. Numerous different types of compound and protein descriptors have been employed for the generation of feature vectors in DTI prediction so far^10,17^, though benchmarking studies have indicated that there is no consensus on what are the sole best compound and target protein descriptors^18,19^. On the compound side, fingerprints are widely used which are binary feature vectors where each dimension represents the presence or absence of sub-structures on a compound. For example, ECFPs^20^ are one of the most widely used fingerprints. To develop DTI prediction methods, a diverse set of ML techniques are employed (together with the feature vectors generated using abovementioned descriptors) such as random forest (RF)^21,22^, support vectors machines (SVM)^22,23^, logistic regression (LR)^24^.

The term “deep learning” (DL) is coined for the novel ML techniques that perform significantly better compared to conventional classifiers especially in the fields of computer vision and natural language processing, mainly due to multiple layers of data abstraction^25^. Deep neural networks (DNN), a group of DL techniques, are artificial neural networks with high complexity, composed of multiple hidden layers^26^. Lately, deep learning based classifiers are also started to be applied for DTI prediction. In one of the earliest applications, Ma *et al*. constructed feed-forward DNN Models using molecular compound descriptors to predict diverse interactions in Merck’s QSAR challenge data sets, and showed that DNNs perform better compared to conventional ML techniques^27^. Lenselink *et al*. proposed a proteochemometric modelling (PCM) based method for DTI prediction, where both compound and target features (i.e., molecular fingerprints for compounds and a custom built composite descriptor - mainly including physicochemical properties-for targets as described by van Westen *et al*.^18^) were employed as 1-D vectors for the training within a multi-layered perceptron DNN architecture^28^. AtomNet, a structure-based virtual screening method, uses convolutional neural networks (CNNs) for drug-target interaction prediction. This method incorporates 3D structural features of known compound-target complexes to model DTIs^29^. Gonczarek *et al*. developed a method that uses specific binding pockets of targets along with fingerprints extracted using the 3-D structural features of compounds^30^. Altae-tran *et al*. proposed a deep learning based method called “iterative refinement long short-term memory” using graph convolutions, where the input of the system is 2D graph structure of compounds^31^. They employed one-shot learning methodology, where the aim is to create predictors for the targets having low number of training samples. Kearnes *et al*. employed graph convolutions to learn features using graph structures of compounds^32^. The field of the deep learning based DTI prediction is still in its infancy and the studies published so far were mostly focused on the applicability of deep learning algorithms and prototyping^27,28,30,32^. The results of these studies have indicated that deep learning has a great potential to advance the field by identifying unknown DTIs at large-scale^28–33^. Apart from the high predictive performance, another advantage of employing deep learning based DTI predictors is the minimal requirement of feature engineering as these algorithms are able to extract complex and meaningful features from the raw data, automatically^34^.

The studies published so far have indicated that DTI prediction is an open problem, where not only novel machine learning algorithms but also new data representation approaches are required to shed light on the un-charted parts of the drug-target interaction space. This effort comprises the identification of novel drug candidate compounds, as well as the repurposing of the existing drugs on the market^35^. Additionally, in order for the DTI prediction methods to be useful in real-world drug discovery and development research (especially non-profit), they should be made available to the research community as tools and/or services via open access repositories. The lack of open access tool/service availability is especially valid for deep learning based DTI predictors.

In this study, we propose DEEPScreen, a convolutional deep neural network based DTI prediction system that utilizes readily available 2-D structural compound images as input features. The main advantages of DEEPScreen is reducing the effort spent on generating complex compound features (i.e., feature engineering) and letting the high-performance deep convolutional neural networks to learn the complex features inherently from the 2-D structural drawings, to produce highly accurate novel drug-target interaction predictions at large scale. Image-based representations of drugs and drug candidate compounds reflect the natural molecular state of these small molecules (i.e., atoms and bonds), which correspond to the features determining their physical interactions with their targets. Recently, image-based or similar structural representations of compounds have been incorporated as input for predictive tasks under different contexts (e.g., toxicity, solubility, and other selected biochemical and physical properties) in the general field of drug discovery and development^36–39^, but have never been investigated in terms of the large-scale prediction of physical interactions between target proteins and drug candidate compounds, which is one of the fundamental steps in early drug discovery. In this work, we aimed to provide such investigation, and as output, we propose a highly-optimised and practical DTI prediction system that covers a significant portion of the known bio-interaction space, with a performance that surpass the state-of-the-art in some tests, and comparable in others.

The proposed system, DEEPScreen, is composed of 704 predictive models, each one is independently optimized to accurately predict interacting small molecule compounds for a unique target protein. DEEPScreen has been validated and tested using various benchmarking datasets, and compared with the state-of-the-art DTI predictors using both conventional and deep ML models.

## Results

### Drug-Target Interaction Prediction with DEEPScreen

In this study, we approached DTI prediction as a binary classification problem. DEEPScreen is a collection of deep convolutional neural networks (CNN), each of which is an individual predictor for a target protein. The system takes drugs or drug candidate compounds in the form of SMILES representations as query, generates 200-by-200 pixel 2-D structural/molecular images using SMILES, run the predictive CNN models on the input 2-D images, and generates binary predictions as active (i.e., interacting) or inactive (i.e., non-interacting) independently for the corresponding target protein (Figure 1). In order to train the target specific predictive models of DEEPScreen with a reliable learning set, manually curated bio-interaction data points are collected from the ChEMBL bioactivity database, and extensively filtered (Figure 2). The technical details regarding both the methodology and the data is given in the Methods section.

**Figure 1.**
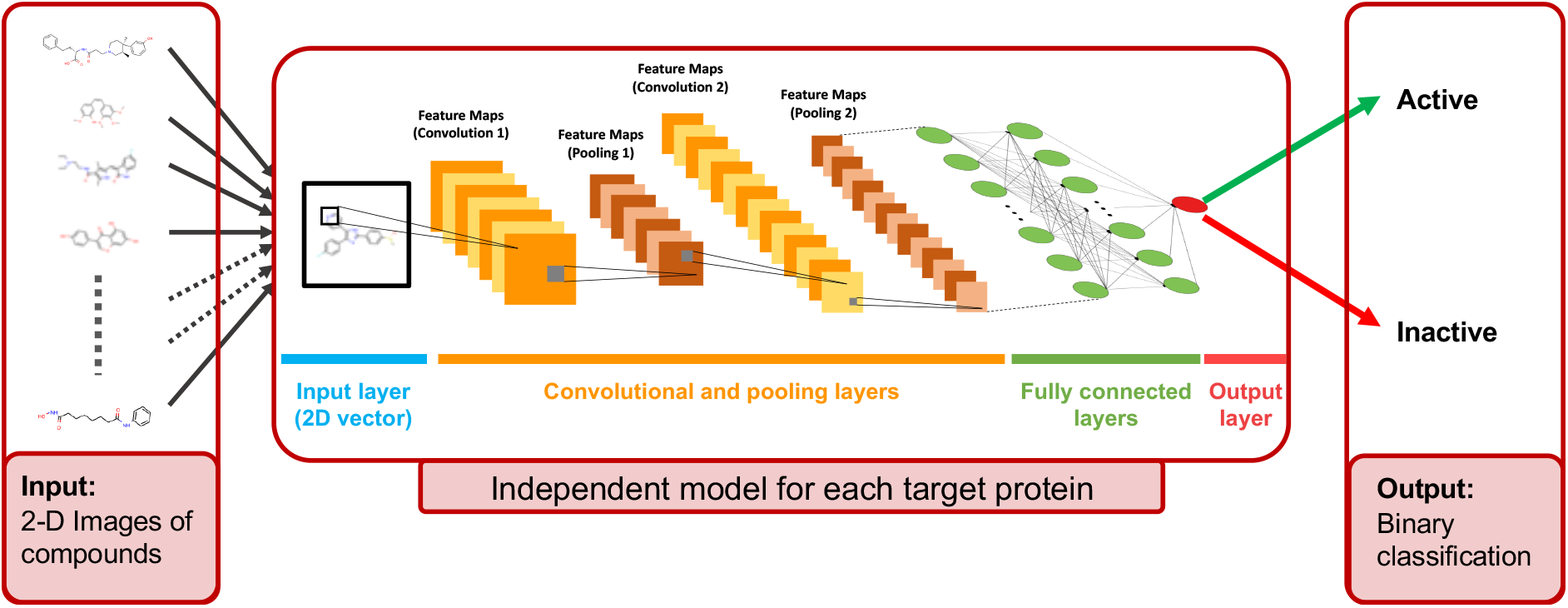
Illustration of the Convolutional Neural Network structure of the proposed DTI prediction system, where the sole input is the 2-D structural images of the drugs and drug candidate compounds (generated from the SMILES representation as a data pre-processing step). Each target protein has an individual prediction model with specifically optimized hyper-parameters (please refer to the Methods chapter). For a query compound, a trained model produces as binary output either as active or inactive for its corresponding target.

**Figure 2.**
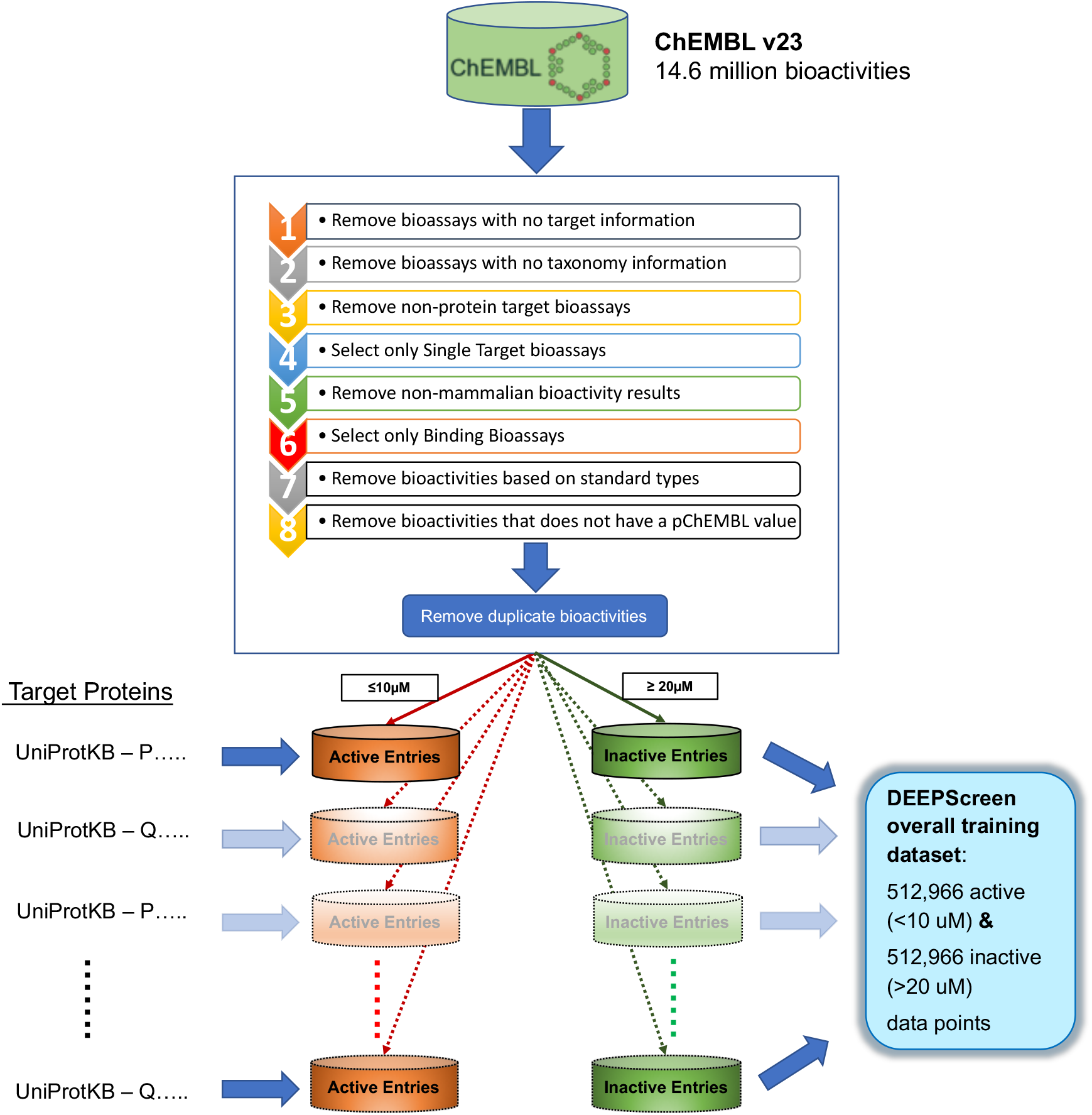
Data filtering and processing steps to create the training dataset of each target protein model. Predictive models were trained for 704 target proteins, each of which have at least 100 known active ligands in ChEMBL database.

### DEEPScreen’s Predictive Performance on ChEMBL Validation and Test Datasets

Hyper-parameter optimization and performance validation of DEEPScreen were accomplished under the same test scheme. For this, several predictive models were trained for each target protein by selecting arbitrary DNN model hyper-parameter values (please refer to the Methods section) and using the corresponding training datasets (i.e., target based interacting and non-interacting compound information) as input. After that, the trained models were run on the validation datasets to obtain the predictive performance (i.e., accuracy, precision, recall, F1-score and MCC), which indicates the effectiveness of the pre-selected hyper-parameters. At the end of the validation procedure, the best performing model (in terms of MCC) of each target was selected, resulting in a total of 704 finalized models. Next, the test performances were calculated by running the finalized models on their corresponding independent test datasets, which have never been used before this point. Figure 3 displays the overall ranked target based predictive performance curves for DEEPscreen (along with other classifiers), with more than 600 of the target protein models (out of 704) received an F1-score higher than 0.8 (average F1-Score and MCC of 0.85 and 0.71, respectively). We also calculated high-level target protein family based average model performances, where results indicated that DEEPScreen performs sufficiently well on all target families (Table 1).

**Figure 3.**
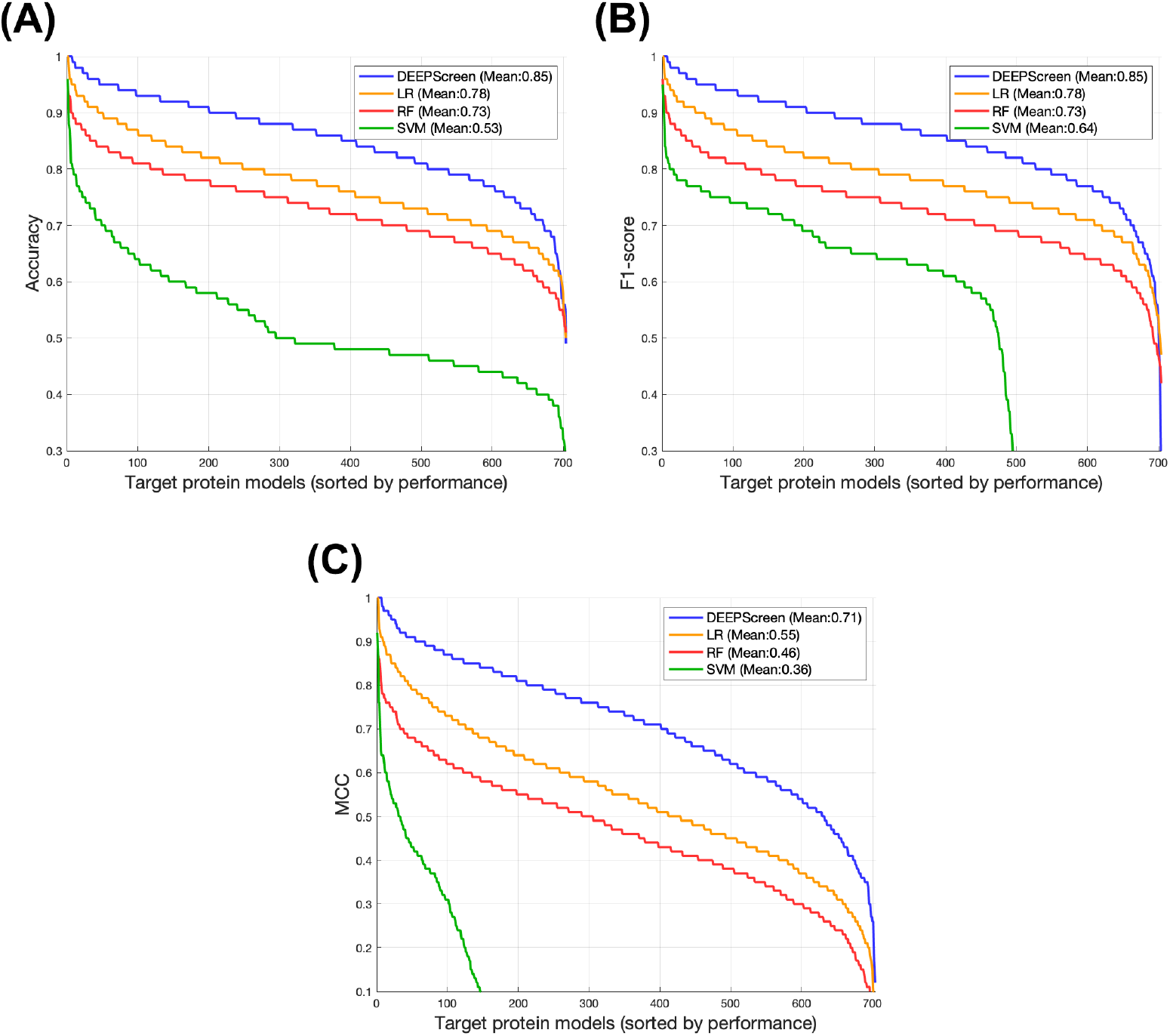
DEEPScreen vs. shallow/conventional classifiers overall predictive performance curves. Each point in the horizontal axis represents a target, the vertical axis represents performance in Accuracy (A), F1-score (B) and MCC (C), respectively. For each classifier, targets are ranked in a descending performance order. Average performance values are given inside the plots.

**Table 1:** Target family-based average predictive performance results of DEEPScreen.

| Family | # of Targets | Average F1-score | Average MCC |
| --- | --- | --- | --- |
| Enzyme | 374 | 0.84 | 0.69 |
| GPCR | 212 | 0.91 | 0.81 |
| Ion Channel | 33 | 0.88 | 0.74 |
| Nuclear Receptor | 27 | 0.89 | 0.77 |
| Other Families | 58 | 0.84 | 0.67 |

### Performance Comparison Between DEEPScreen and Conventional/Shallow ML Classifiers

We compared the performance of DEEPScreen against conventional classifiers that are frequently employed in DTI prediction (e.g., random forest, SVM and logistic regression) using the exact same training/test sets and input features. These conventional classifiers generally accept 1-D (column type) feature vectors, therefore, we flattened our 200-by-200 images to be used as input to all of the conventional classifiers. Thus, the performance comparison solely reflects the gain of employing deep convolutional networks as opposed to conventional classifiers. The models parameters of the shallow classifiers were optimized on the validation dataset and the finalized performances were measured using the independent test set, similar to the evaluation of DEEPScreen. Figure 3 displays the overall results in accuracy, F1-score and MCC, respectively; where DEEPScreen performed significantly better compared to all conventional classifiers employed in the test. According to our results, the sole best classifier was DEEPScreen for 642 targets (LR for 42, RF for 4, SVM for 2 targets), out of the total 704. Figure 4 shows target protein based predictive performance (in terms of MCC) z-score heatmap for DEEPScreen and the shallow classifiers, where each horizontal block corresponds to a target family. Considering the results in Figure 4, DEEPScreen performed significantly better for all families, LR was the second (LR’s enzyme family performance was better compared to other protein families), RF and SVM came at the last two places. Overall, the target models with low performance (MCC < 0.5) were mostly belong to the enzyme family. An interesting observation here is that, except from a few target models from each protein family where multiple classifiers performed well, DEEPScreen and LR models display opposite trends in predictive performance. Despite being a widely used conventional classifier in DTI prediction, RF performed poorly, compared to both DEEPScreen and LR, when employed with image-based features. For most of the cases where LR was the best classifier, DEEPScreen came second and RF the third. There was no significant difference between the protein families in terms of the classifiers rankings, though, DEEPScreen’s superiority was more pronounced on the families of nuclear receptor, ion channel and others. Finally, SVM was unable to learn the data in many target models, and classify all instances into just one class.

**Figure 4.**
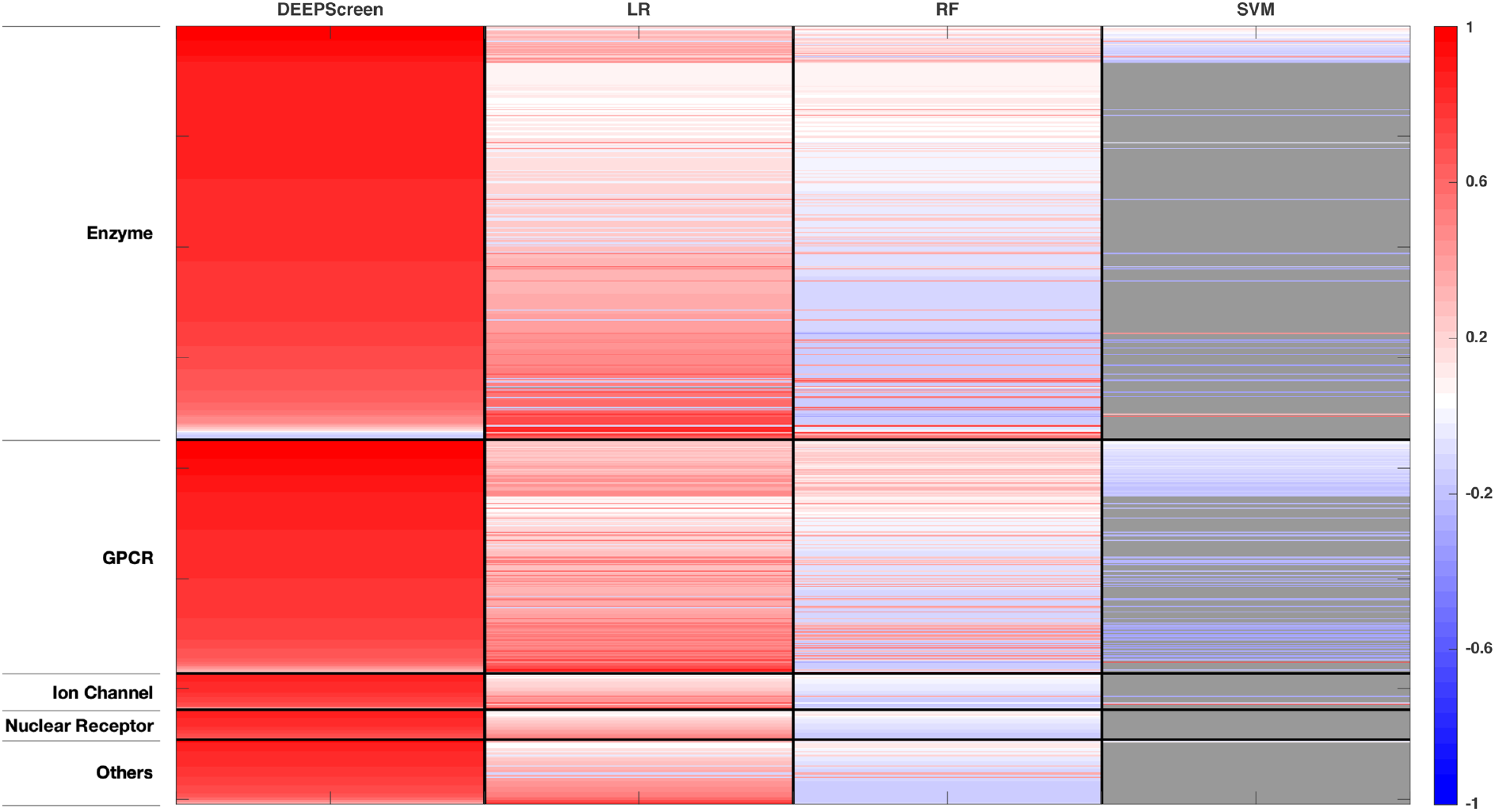
Target-based (rows) maximum predictive performance (MCC-based) heatmap for DEEPScreen and shallow classifiers (columns) (LR: logistic regression, RF: random forest, SVM: support vector machine). For each target protein (row), classifier performances are shown in the shades of red (i.e., high performance) and blue (i.e., low performance) colours according to Z-scores, calculated individually for each target model (row). Rows are arranged in blocks according to target families. The length of a block is proportional to the number of targets in its corresponding family (enzymes: 374, GPCRs: 212, ion channels: 33, nuclear receptors: 27, others: 58). Within each block, targets are arranged according to descending performance from top down with respect to DEEPScreen. Grey colour signifies the models where learning was not possible.

### State-of-the-art DNN-based Methods Performance Comparison

We compared the results of DEEPScreen with five other deep learning based DTI prediction methods that represent the current state-of-the-art (please refer to the Introduction section) by employing the same datasets used in the corresponding studies. For this analysis, we re-trained and tested DEEPScreen using the exact same experimental settings and evaluation metrics that were described in the respective articles^28–32^. A total of 3 different benchmark datasets were employed for this purpose (please refer to the Methods section). Two of these datasets (i.e., MUV and DUD-E) are frequently employed in DTI prediction studies and the performance results of DEEPScreen on these datasets are directly comparable with any study in the literature, where the same benchmark sets are used. The results of this analysis reflect both the effectiveness of employing 2-D images of compounds as the input and the constructed CNN-based architecture. Table 2 shows the results of DEEPScreen along with the state-of-the-art performances reported in the respective articles. As shown, DEEPScreen performed significantly better compared to one method, comparable to another two, and its score is slightly lower than the methods that employ 3-D structural information. It is important to note that the methods employing 3-D structural features of the target proteins may provide better representations to model DTIs; however, they are highly computationally intensive. Also, 3-D structural information (especially the target-ligand complexes) is only available for a small portion of the DTI space, as a result, their coverage is low.

**Table 2:** The average predictive performance comparison between DEEPScreen and various state-of-the-art deep learning based DTI predictors.

| Dataset | Method | Architecture | Performance (Metric) |
| --- | --- | --- | --- |
| ChEMBL Temporal-split Dataset | DEEPScreen | ConvNets with 2-D Images | 0.45 (MCC) |
|  | Lenselink *et al.* | Feed-forward DNN PCM (best model) | 0.33 (MCC) |
| Directory of Useful Decoys -Enhanced (DUD-E) Dataset | DEEPScreen | ConvNets with Images | 0.83 (AUROC) |
|  | AtomNet | 3D Structural Convolutional NNs | 0.85 (AUROC) |
|  | Gonczarek *et al.* | Feed-forward multi-task DNNs | 0.90 (AUROC) |
| Maximum Unbiased Validation (MUV) Dataset | DEEPScreen | ConvNet with Images | 0.74 (AUROC) |
|  | Kearnes *et al.* | Graph convolution NNs | 0.75 (AUROC) |
|  | Altae-Tran *et al.* | One shot learning + graph conv. | 0.66 (AUROC) |

## Discussion

In this work, we have shown that deep convolutional neural networks can be utilized to successfully predict the drug-target interactions, using only the 2-D structural images of drugs and drug candidate compounds. The proposed method, DEEPScreen, has been tested on various benchmarking datasets and compared with both conventional and the state-of-the-art DTI prediction methodologies to reveal that it performs well.

In DEEPScreen, we modelled the interactive properties of each target protein independently in a separate convolutional network. This allowed the target based optimization of hyper-parameters, as well as the regular models parameters. In most of the ML method development studies, hyper-parameters are arbitrarily pre-selected without further optimization (especially when there are high number of models as in DEEPScreen), due to extremely high computational burden. However, hyper-parameters are an important part of the model architecture and significantly contribute to the predictive performance. Using the hyper-parameter value alternatives given in Table 3 (and considering their combinations with each other), we evaluated hundreds of models for each target on average, resulting in more than 100,000 model training and evaluation jobs in total. The main advantage of this approach is the elevated predictive performance, which was indicated by the results of the performance comparison tests. One important concern in ML method development is the problem of overfitting. We employed the neuron drop-out technique in order to prevent overfitting, which is a widely accepted approach for DNN training. The results of the independent tests confirmed that overfitting was not a problem for DEEPScreen.

**Table 3:** Hyper-parameter types and their values that were tested during the training of the DEEPScreen system.

| Hyper-parameter Name | Range |
| --- | --- |
| Input Normalization | Yes |
|  | No |
| Learning rate | 0.0005 |
|  | 0.0001 |
|  | 0.005 |
|  | 0.001 |
|  | 0.01 |
| Number of neurons in the fully-connected layers | NA* |
| Optimizer | Adam (default) |
|  | Momentum (default) |
|  | RMSprop (default) |
| Mini-batch size | 32 |
|  | 64 |
| Drop-out rate | 0.5 |
|  | 0.6 |
|  | 0.8 |
| Batch Normalization | Yes |

\* There are two fully-connected layers after the convolutional layers for the AlexNET model. The number of neurons that were tested these layers were (128,16), (256,128), (512,32), (1024,32), (2048,2048). For the in-house convolutional neural network architecture, there were one fully connected-layer and the number neurons tested for the corresponding layer was 16, 32, 128, 256, 512.

One of the critical points in the computer vision tasks is the system robustness concerning the differences in the representations of the object of interest, such as the viewing angle or the scale. In DEEPScreen, input images are standardized by computationally generating them using SMILES representations, this way all images have similar representations in terms of rotation and scaling. In DEEPScreen, we selected a considerably loose threshold (i.e., 10 uM) bioactivity value to label training instances as active (i.e., interacting), which in turn resulted in high number of predictions. The true experimental bioactivity measurement of a pair that is accurately predicted as active can go as high as 10 uM, which can be considered extremely high especially for certain target families (e.g., kinases and GPCRs). However, our aim in developing DEEPScreen was to aid experimental researchers in drug discovery and repurposing by providing all matches that would potentially raise an interest. We believe that the methodology proposed here can be employed to produce large-scale production of novel drug-target interactions, which can be utilized by the experimental and computational researchers to aid their work on drug design and repurposing.

## Methods

### Generation of the Training Datasets

ChEMBL database (version 23) was employed to create the training dataset of DEEPScreen. There are 14,675,320 data points (i.e., bioactivity measurements) in ChEMBL v23. We applied several filtering and pre-processing steps on this data to create a reliable training dataset. First of all, data points were filtered with respect to “target type” (i.e., single protein), “taxonomy” (i.e., human and selected model organisms), “assay type” (i.e., binding and functional assays) and “standard type” (i.e., IC50, XC50, EC50, AC50, Ki, Kd and Potency) attributes, which reduced the set to 3,919,275 data points. We observed that there were duplicate measurements inside this dataset that are coming from different bioassays (i.e., 879,848 of the bioactivity data points belonged to 374,024 unique drug-target pairs,). To handle these cases, we identified the median bioactivity value for each pair and assigned this value as the sole bioactivity measurement. At the end of this application, 3,413,451 bioactivity measurements were left. This dataset contained data points from both binding and functional assays. In order to eliminate a potential ambiguity considering the physical binding of the compounds to their targets, we discarded the functional assays and kept the binding assays with an additional filtering on “assay type”. Finally, we removed the bioactivity measurements without any pChEMBL value, which is a standardized value to obtain comparable measures of half-maximal response (e.g., IC50, EC50, Ki, Kd and Potency) on a negative logarithmic scale. The presence of a pChEMBL value for an activity point indicates that the corresponding record has been curated and thus reliable. After these processing steps, the number of bioactivity points were 769,935.

Subsequently, we constructed positive (active) and negative (inactive) training datasets as follows: For each target, compounds with bioactivity values <= 10 μM were selected as positive training samples and compounds with bioactivity values >= 20 μM were selected as negative samples. In DEEPScreen only the target proteins with at least 100 active ligands we modelled in order not to lose the statistical power. This application provided models for 704 target proteins from multiple highly studied organisms. These organisms, together with the distribution of target proteins for each organism are: Homo sapiens (human): 523, Rattus norvegicus (rat): 88, Mus musculus (mouse): 34, Bos taurus (Bovine): 22, Cavia porcellus (Guinea pig): 13, Sus scrofa (Pig): 9, Oryctolagus cuniculus (Rabbit): 5, Canis familiaris (dog): 3, Equus caballus (horse): 2, Ovis aries (Sheep): 2, Cricetulus griseus (Chinese hamster): 1, Mesocricetus auratus (Golden hamster): 1 and Macaca mulatta (Rhesus macaque): 1. The UniProt accessions, encoding gene names, ChEMBL ids and the taxonomic information of these proteins are given in the repository of DEEPScreen. Each target’s training set contained a mixture of activity measurements with roughly comparable standard types (e.g., IC50, EC50, Ki, Kd and Potency).

The selection procedure explained above generated positive and negative training datasets with varying sizes for each target. In order to balance these datasets, we selected negative samples equal to the number of positive instances. However, for many targets, the number of negative points were lower than the positives. In these cases, we applied a target similarity-based inactive dataset enrichment method to populate the negative training sets (instead of randomly selecting compounds), using the idea of similar targets have similar actives/inactives. For this, we first calculated pairwise similarities between all target proteins within a BLAST search. For each target having insufficient number of inactive compounds, we sorted all remaining target proteins with descending sequence similarity. Then, starting from the top of the list, we populated the inactive dataset of the corresponding target using the known inactive compounds of similar targets, until the active and inactive datasets are balanced. We applied 20% sequence similarity threshold, meaning that we did not consider the inactives of targets, whose sequence similarity to the query protein is less than 20%. The finalized training dataset for 704 target proteins contained 512,966 active data points (< 10 uM) and the same number of inactive data points (> 20 uM). Before the negative dataset enrichment procedure, the total number of inactive instances for 704 targets were 35,567. Both the pre-processed ChEMBL dataset (769,935 data points) and the finalized active and inactive training datasets for 704 targets are given in the repository of DEEPScreen. We believe the resulting bioactivity dataset is reliable, and it can be used as a standard training/test sets in future DTI prediction studies. The training data filtering and pre-processing operations are represented in Figure 2.

For each target protein model, 80% of the training samples (from the positives and the negatives datasets) were randomly selected for training/validation dataset, and the remaining 20% was reserved for later use in the independent performance test procedure. Also, 80% of the training/validation dataset was employed for system training and 20% of this dataset was used for validation, during which the hyper-parameters of the models were optimized.

### Representation of Input Samples and the Generation of Feature Vectors

In DEEPScreen system, each compound is represented by a 200-by-200 pixel 2-D image displaying the molecular structure (i.e., skeletal formula). Although 2-D compound images are readily available in different chemical and bioactivity databases, there is no standardization in terms of the representation of atoms/bonds, functional groups and the stereochemistry. Due to this reason, we employed SMILES strings of compounds to generate the 2-D structural images, since SMILES is a standard representation that can be found in the open access bioactivity data repositories, which contain all of the necessary information to generate the 2-D images. We employed the RDkit tool Python package (v2016.09.4) for the image generation^40^. A few examples from the generated images are shown in Figure 1.

### Architecture of the DEEPScreen System

Convolutional neural networks (CNN) are a specialized group of artificial neural networks consisting of alternating, convolution and pooling layers, which extracts features automatically^25,41^. They run a small window over the input feature vector at both training and testing phases as a feature detector and learn various features from the input regardless of their absolute position within the input feature vector. Convolution layers compute the dot product between the entries of the filter and the input, producing an activation map of that filter. Whereas, pooling layers combine the outputs of neuron clusters at one layer into a single neuron in the next layer (i.e., a non-linear down-sampling operation). Although the most standard from of CNNs employ 2-D convolutions, 1-D or 3-D convolutions are applied as well. The CNNs have been dominating image processing area in the last few years, achieving significantly higher performances compared to the state-of-the-art of the time^25,42,43^.

In this study, we considered the DTI prediction as a binary classification problem, where the output can either be positive (i.e., active, interacting or “1”) or negative (i.e., inactive, non-interacting or “0”), referring to the relation between the query compound and the modelled target protein. For this purpose, an individual model was created for each target protein (i.e., the single task approach). In terms of the employed CNN architectures, we chose 3 options: ImageNET^42^, AlexNET^43^; and an in-house built CNN composed of 5 convolutional + pooling and 1 fully-connected layer preceding the output layer. A generic representation of the constructed CNN models is given in Figure 1.

Subsequently, both positive (i.e., active) and negative (i.e., inactive) training datasets were prepared for each target protein according to the rules explained in the training dataset generation sub-section. Also, the input feature vectors (i.e., 200-by-200 pixel 2-D images) of the compounds in our dataset were automatically constructed via RDkit. For each model, we carried out comprehensive hyper-parameter optimization tests. The list of the hyper-parameters and the value selections are given in Table 3.

### Independent Test Datasets for The Comparison with The State-of-the-art

The first dataset was obtained from the study by Lenselink *et al*.^28^. In this study, the authors created a high quality ChEMBL (v.20) bioactivity dataset that includes 314,767 bioactivity measurements corresponding to single protein targets with at least 30 data points. They used pChEMBL = 6.5 (roughly 300 nM) bioactivity value threshold to create active and inactive compound datasets for each target. The authors evaluated their method with a test dataset created by a temporal split, where for each target protein, all the bioactivity data points reported in the literature prior to 2013 were used in the training, and the newer data points were gathered for the test dataset. This test dataset is more challenging for ML classifiers compared to any random-split dataset.

The second independent test dataset employed in this study was DUD-E, a well-known benchmarking set for DTI prediction, which includes curated active and inactive compounds for 102 targets. The active compounds for each target was selected by first clustering all active compounds based on the scaffold similarity and selecting representative actives from each cluster. The inactive compounds were selected to be similar to the active compounds in terms of the physicochemical descriptors, but dissimilar considering the 2-D fingerprints^44^. The benchmark dataset consists of 102 targets, 22,886 actives (an average of 224 actives per target) and 50 property-matched decoys for each active, which were obtained from the ZINC database^44^.

The third and final test dataset was Maximum Unbiased Validation (MUV), another widely-used benchmark set, composed of active and inactive (decoy) compounds for 17 targets^45^. MUV dataset was generated from the PubChem Bioassay database. The active compounds in this dataset was selected to be structurally different from each other, therefore, it is a challenging benchmark dataset, which avoids the bias rooting from highly similar compounds ending up in both train and test splits. There are 17 targets in MUV dataset, together with 30 actives and 15000 decoys for each target.

### Performance Evaluation Metrics

To evaluate the predictive performance of DEEPScreen and to compare our results with other DTI prediction methods we mainly used 3 evaluation metrics, which are F1-score, Matthews correlation coefficient (MCC) and area under receiver operating characteristic curve (AUROC). The formulas of these evaluation metrics are given below together with precision and recall that make up F1-score:

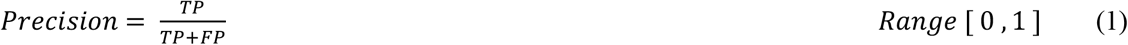

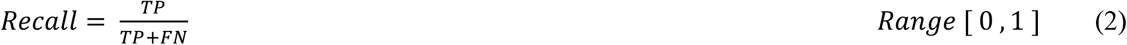

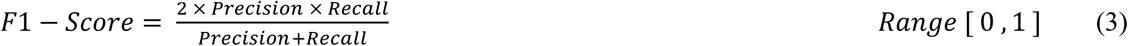

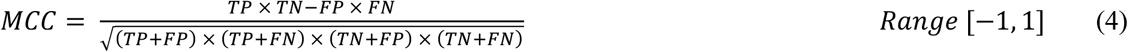

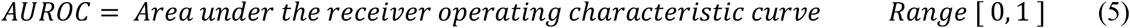

In the above equations, TP (i.e., true positive) represents the number of correctly predicted interacting compound-target pairs, FN (i.e., false negative) represents the number of interacting compound-target pairs, that are predicted as non-interacting (i.e., inactive). TN (i.e., true negative) denotes the number of correctly predicted non-interacting compound-target pairs, whereas FP (i.e., false positive) represents the number of non-interacting compound target pairs, which are predicted as interacting.

